# 40S Ribosome Remodeling Triggers 18S rRNA U-tailing and DIS3L2-dependent Decay

**DOI:** 10.64898/2026.09.11.750429

**Authors:** Akruti Shah, Aaztli R. Coria, Jennifer T. Miller, Emilien Orgebin, Wilfried Guiblet, Acong Yang, Shuo Gu, Colin Chih-Chien Wu

**Affiliations:** Center for Cancer Research, National Cancer Institute, National Institutes of Health, Frederick MD 21702, USA; Advanced Biomedical Computational Science, Frederick National Laboratory for Cancer Research, Frederick, MD 21702, USA

## Abstract

Eliminating defective ribosomes through quality control is essential for accurate protein synthesis. However, the mechanisms that commit ribosomal subunits to decay remain poorly defined. Here, we identify a tandem mechanism in which ubiquitin-mediated ribosome remodeling and 18S rRNA uridylation lead to 40S ribosomal subunit decay. Specifically, we use an *in vitro* reconstitution system to show that the atypical kinase RIOK3 displaces the ribosomal protein eS26 from 40S subunits, exposing the 3’ end of 18S rRNA. Nanopore direct RNA sequencing and motif analyses reveal that this remodeling event promotes oligo-uridylation, generating uridylated 18S rRNA decay intermediates. We further show that this uridylated 18S rRNA is selectively degraded by the 3’-5’ exoribonuclease DIS3L2. Moreover, DIS3L2-mediated exoribonucleolytic cleavage triggers endoribonucleolytic decay of the 18S rRNA, amplifying turnover. Together, our findings define a stepwise mechanism in which ribosome remodeling and RNA tailing commit defective 40S subunits to elimination, establishing a mechanistic framework for ribosome surveillance in mammalian cells.

## INTRODUCTION

Ribosomes are among the most abundant macromolecular complexes in the cell and play a central role in protein synthesis. Yet, they are also highly sensitive to physiological stress, functioning as regulatory hubs that sense translational dysfunction and engage quality control pathways to restore cellular homeostasis ^1,2^. In this context, ubiquitination of ribosomal proteins has emerged as a key signal of translational perturbations, marking aberrant ribosomes for downstream turnover and surveillance pathways ^1,3^. Recent studies have shown that diverse perturbations—including nonfunctional 18S rRNA, amino acid starvation, and disruption of 60S ribosome biogenesis— trigger ubiquitination of 40S ribosomal proteins and promote selective degradation of the small ribosomal subunit ^4–7^. Despite these advances, a key mechanistic gap remains unresolved: how are ubiquitinated 40S ribosomes committed to destruction?

One molecular pathway that mediates this response is 18S nonfunctional rRNA decay (NRD) ^8,9^, a surveillance mechanism that selectively eliminates defective 40S subunits containing dysfunctional 18S rRNA. We and others showed that a single-nucleotide mutation in the decoding center of 18S rRNA (A1824C in human and A1755C in budding yeast) stalls ribosomes at translation initiation sites, thereby triggering decay through ubiquitination of ribosomal proteins uS3 and uS5^5,10^. Moreover, we and others identified RIOK3 as a downstream effector that binds these ubiquitinated 40S subunits to promote 18S rRNA decay ^5,11,12^. Importantly, similar mechanisms operate in response to other stress conditions that impair productive translation, including amino acid starvation and defects in 60S ribosomal subunit biogenesis ^7,11,12^, supporting a model in which ubiquitin mark is a general prelude to 40S subunit turnover. Additionally, structural analysis of 40S decay intermediates further indicates that degradation begins near the 3ʹ end of 18S rRNA, suggesting that defective 40S subunits may undergo localized remodeling to facilitate decay ^11^. Despite diverse translational stress conditions converging on 18S rRNA degradation, the mechanism underlying decay remains unclear.

Among eleven terminal nucleotidyltransferases, TUT4 and TUT7 are the two well-established terminal uridylyltransferases (TUTases) for modifying cytoplasmic structured noncoding RNAs.

Oligo-uridine tailing can function as a degradation signal for RNA turnover in multiple surveillance pathways ^13–18^. Indeed, U-tailing promotes recognition by the cytoplasmic 3ʹ-5ʹ exoribonuclease DIS3L2, which preferentially binds and degrades uridylated RNA substrates ^13–15,19,20^. In addition to its established roles in the turnover of pre- and mature microRNAs ^21–23^, DIS3L2 has also been linked to degradation of uridylated 5.8S rRNA biogenesis intermediates ^24^.

Here, we show that RIOK3 promotes the release of ribosomal protein eS26 from ubiquitinated 40S subunits, thereby exposing the 3ʹ end of 18S rRNA and enabling subsequent uridylation of the accessible 3’ terminus. We further show that TUTases, TUT4 and TUT7, are in part responsible for this modification. The 3’-5’ exoribonuclease DIS3L2 then initiates degradation from the 3’ terminus of 18S rRNA, promoting endoribonucleolytic cleavage events. Importantly, these cleavage events further amplify 40S subunit turnover by generating additional substrates for uridylation-dependent DIS3L2-mediated decay. Together, our findings establish that defective mammalian 40S subunits are eliminated through a uridylation-dependent decay mechanism, revealing how ribosome remodeling, terminal RNA tailing, and exoribonucleolytic degradation are coupled during ribosome quality control.

## RESULTS

### RIOK3 releases eS26 to promote 18S rRNA decay

In light of prior studies implicating the release of ribosomal protein eS26 (RPS26) in ribosome repair ^25,26^, and since eS26 is located near the 3’ terminus of 18S rRNA in the 40S ribosomal subunit^27^, we deduced that RIOK3 might displace eS26 to expose the 3’ end of 18S rRNA, thereby rendering it accessible to exoribonucleases. To test this hypothesis, we reconstituted RIOK3 activity *in vitro* with purified components. We performed *in vitro* ubiquitination with purified 40S ribosomal subunits containing HA-tagged uS3 isolated from USP10/RIOK3 double-knockout (DKO) cells, thereby eliminating potential contamination by endogenous RIOK3 and USP10, a deubiquitinase that acts on ubiquitinated uS3 and uS5 ^28^. After immobilization of 40S subunits on α-HA beads, uS3 and uS5 ubiquitination was carried out using recombinant ubiquitin, RNF10 (E3), UbcH5c (E2), and Ube1 (E1) (Figures S1A-S1B). Following the ubiquitination reaction, RNF10, UbcH5c, and Ube1 were removed by stringent washes. The ubiquitinated 40S subunits were then directly incubated with recombinant wild-type (WT) or mutant RIOK3 (Figure S1C), thereby reconstituting 40S turnover events (Figure 1A).

**Figure 1.**
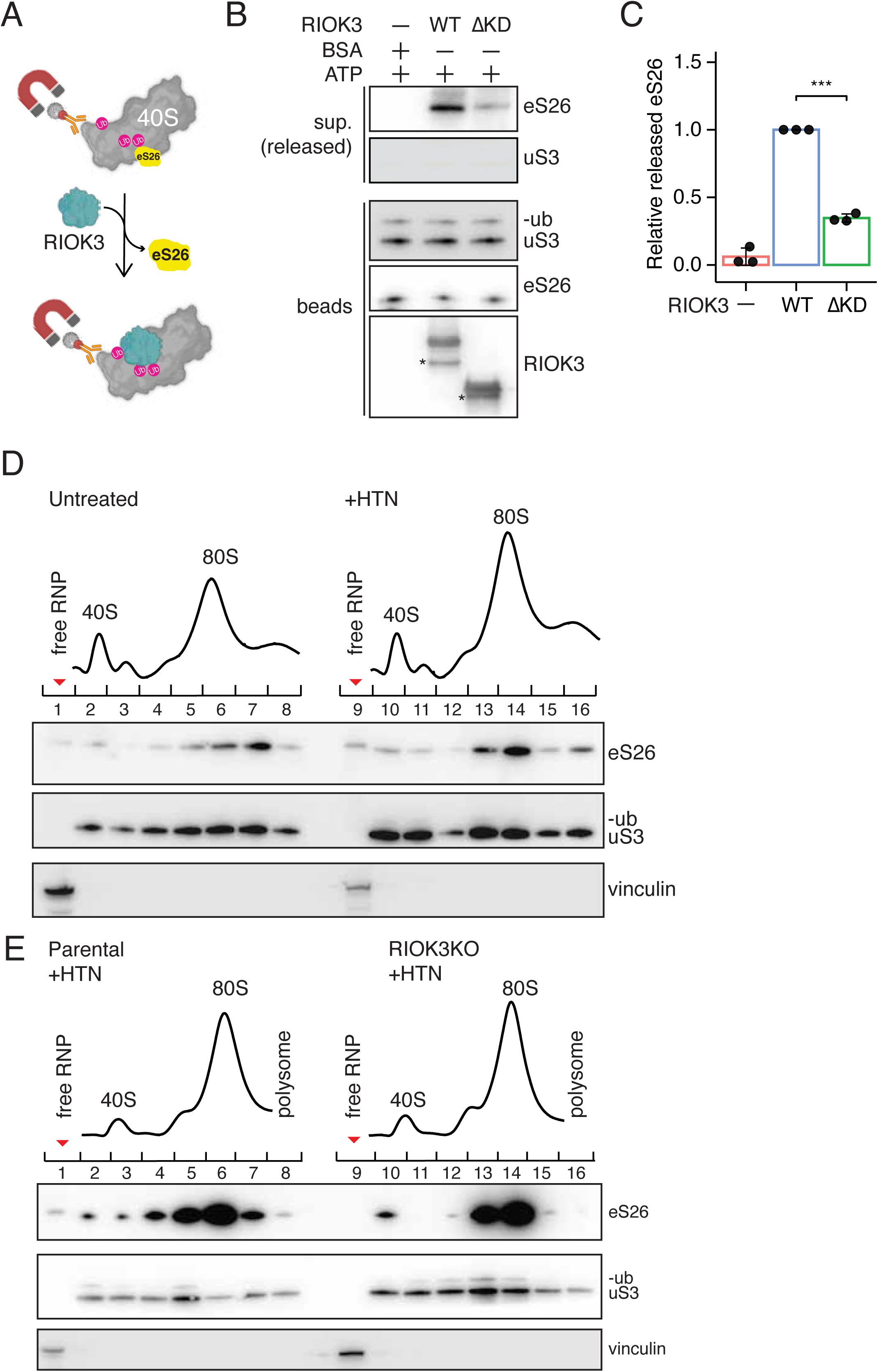
RIOK3 displaces ribosomal protein eS26 during 40S subunit decay. (A) Schematic depicting *in vitro* displacement assays of eS26 by RIOK3. 40S subunits containing HA-tagged uS3 were immobilized on magnetic beads and recombinant RIOK3 was added to reactions. Released eS26 is detected by its presence in the supernatant. (B) Representative immunoblotting of eS26 release reactions in the presence of BSA, RIOK3 WT or RIOK3 ΔKD recombinant proteins. Released (supernatant, sup) and immobilized (beads) fractions were analyzed with indicated antibodies. uS3 serves as controls. (C) Quantification of relative levels of eS26 release shown in (B), (n= 3). (D) Representative polysome profiles from 10-30% sucrose gradients from untreated (left) and HTN-treated (right) HEK293T cells. Fractions were analyzed by immunoblotting with indicated antibodies, (n= 2). Vinculin, which migrates predominantly in free RNP fractions, serves as a loading control. Red arrowheads indicate free RNP fractions. Comparison of lanes 1 and 9 indicate the release of eS26 following HTN treatment. (E) Similar to (D), representative polysome profiles and immunoblotting for HTN-treated H293T parental (left) and RIOK3 KO cells (right), (n= 3).

Addition of RIOK3 WT promoted the release of eS26 from immobilized 40S subunits, leading to significant enrichment in the supernatant (sup) fraction compared to the BSA control (Figures 1B and 1C). In contrast, a distinct ribosomal subunit, ubiquitinated uS3, remained associated with immobilized 40S subunits (Figure 1B). We also tested RIOK3 ΔKD, which is critical for 18S NRD (Figures S1D-S1E) and 40S subunit turnover under stress ^11,12^. Importantly, RIOK3 ΔKD released eS26 less efficiently than full-length RIOK3, although RIOK3 ΔKD bound to ubiquitinated 40S subunits with comparable efficiency (Figure S1F). Furthermore, WT RIOK3 was unable to release eS26 in the absence of ATP (Figure S1G). Together, these results indicate that RIOK3 promotes the dissociation of eS26 from ubiquitinated 40S subunits in an ATP-dependent manner and that its kinase domain is required.

We next tested RIOK3-mediated eS26 release in cells. Prior studies showed harringtonine (HTN) treatment induces ribosome stalling at translation initiation sites ^29^, resulting in ubiquitination of uS3 and uS5 ^6,7,30^ as well as mimicking the defect in the 18S decoding center ^5^. We therefore utilized HTN treatment to trigger 40S subunit degradation and monitored eS26 release by polysome profiling followed by immunoblotting. Upon HTN treatment, eS26 was detected in the free RNP fraction (Figure 1D, compare lane 1 to 9) compared to the untreated control, indicating its dissociation form 40S subunits. Importantly, this release of eS26 was abolished in RIOK3 KO cells (Figure 1E, compare lane 1 to 9). These results demonstrate RIOK3-dependent dissociation of eS26 from ubiquitinated 40S subunits in cells.

### 18S rRNA decay intermediates accumulate uridylated tails

18S rRNA degradation is initiated with successive endo- and exoribonucleolytic cleavage events ^5,11^. To further dissect 18S rRNA decay intermediates, we performed Nanopore direct RNA sequencing on RIOK3-bound 18S rRNA from HEK293T cells treated with HTN. Reads were aligned to the 18S rDNA reference sequence and untemplated nucleotides were extracted for downstream analysis (Figure 2A). To improve resolution, given the lower accuracy of Nanopore sequencing compared to Illumina, we additionally employed an amplicon-based sequencing approach coupled to Illumina sequencing (Figure S2A) to characterize these untemplated nucleotides. Motif analysis of these untemplated reads revealed a striking and unexpected enrichment of uridine residues, which generates a strong oligo-uridine signature (Figure 2B).

**Figure 2.**
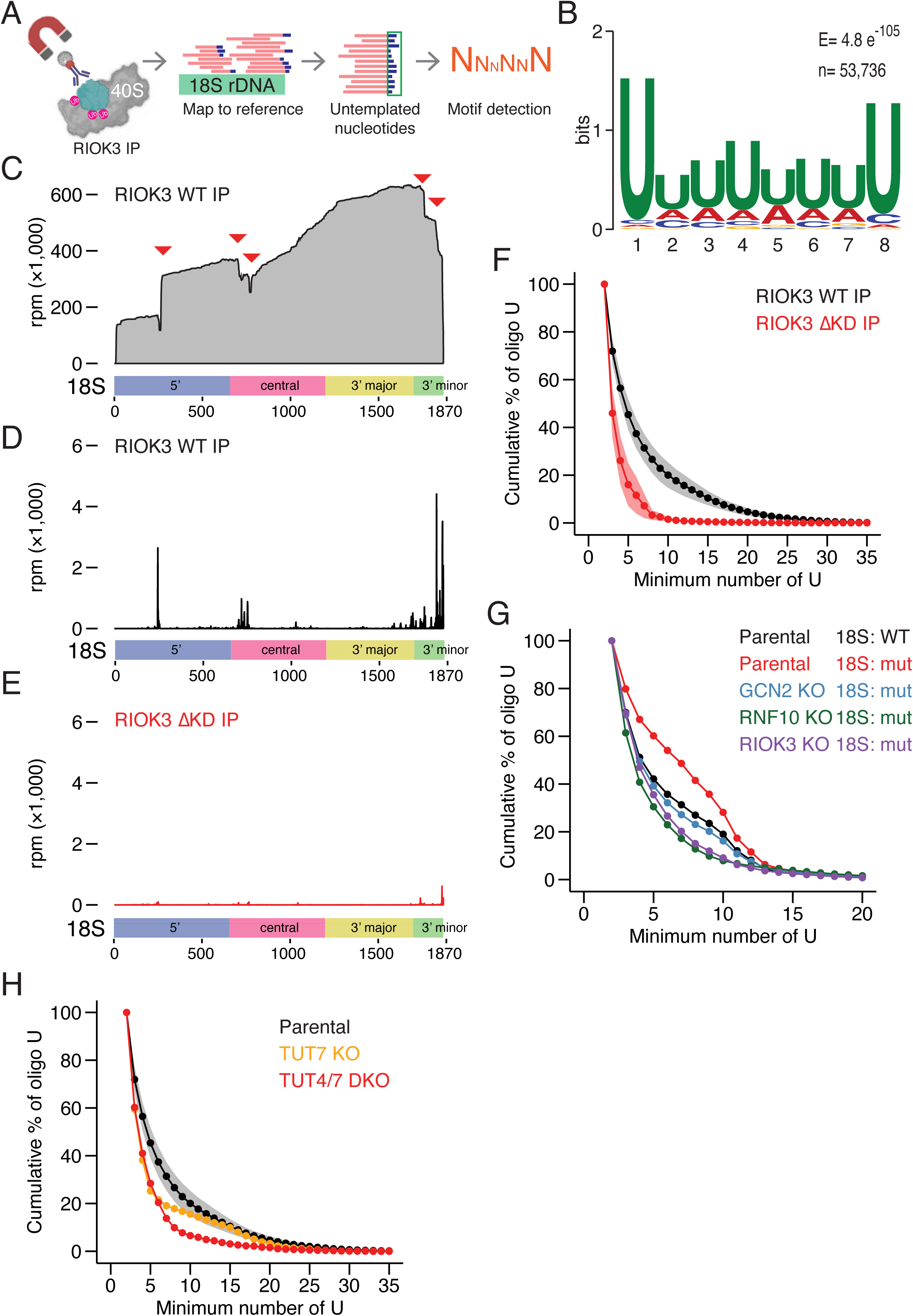
RIOK3-bound 18S rRNA decay intermediates are uridylated at their 3’ termini. (A) Schematic of the untemplated nucleotide analytical pipeline for 18S rRNA decay intermediates. See also Figure S2A. (B) MEME motif analysis revealed oligo-uridine motif enriched in untemplated nucleotides at the 3’ ends of 18S rRNA decay intermediates. E represents the E-value significance; n denotes the number of contributing sites out of 53,736 input sequences. (C) Representative gene model of 18S rRNA depicting the read coverage of Nanopore direct RNA sequencing reads of RIOK3-bound 18S rRNA decay intermediates isolated from HTN-treated HEK293T cells, (n= 3). Positions of increased or decreased read coverage are indicated by red arrowheads. Bottom: Schematic representation of 18S rRNA domain organization (color-coded). (D-E) Representative gene models of 18S rRNA illustrating the 3ʹ ends of oligo-uridylated (U≥ 2) from RIOK3 WT (D) and RIOK3 ΔKD (E) immunoprecipitation (IP), (n≥ 2). Y-axis represents read density in reads per million (rpm x1,000), indicating the relative abundance of uridylated decay intermediates across the 18S rRNA. (F) Cumulative percentage of uridylated reads across the 18S rRNA from RIOK3 WT (black) and RIOK3 ΔKD (red) IP isolated from parental HEK293T cells treated with HTN (3 µg/mL), (n≥ 2). (G) Cumulative percentage of uridylated reads across the 18S rRNA for 18S:WT reporter isolated from parental cells (black) and 18S:A1824C reporter (mut) isolated from parental (red), GCN2 KO (blue), RNF10 KO (green), and RIOK3 KO cells (purple). Data were obtained from SRA (accession number: PRJNA1195343) ^5^. (H) Cumulative percentage of uridylated reads across the 18S rRNA from RIOK3-bound 18S rRNA decay intermediates isolated from HTN-treated HEK293T parental (black), TUT7 KO (orange), and TUT4/7 DKO (red) cells.

Read coverage across RIOK3-bound 18S rRNA, which reflects the sequencing depth, revealed degradation in the 5’, central, and 3’ minor domains (Figure 2C), consistent with successive endo- and exoribonucleolytic cleavages during decay ^5,11^. Mapping of the 3’ ends of uridylated reads (defined as reads containing at least two untemplated uridines) across the 18S rRNA showed that these reads coincided well with the cleavage events and were particularly enriched toward the 3ʹ terminus (Figure 2D), suggesting that 18S rRNA decay intermediates are uridylated after endo- and exoribonucleolytic cleavage. Though there has been much interest in RNA U-tailing ^13–18^, its role in the degradation of mature 40S ribosomes has remained unrecognized, likely because the 18S rRNA is structurally sequestered within the 40S subunit.

Notably, ∼4% of RIOK3-bound 18S rRNAs were uridylated, with a broad distribution extending up to 30 nucleotides, compared to input 18S rRNA reads (Figure S2B). Consistently, re-analysis of data from prolonged amino acid starvation ^11^ revealed the same uridylation trend in RIOK3-bound 18S rRNA decay intermediates (Figure S2C). This striking enrichment in the RIOK3-bound fractions underscores its role in promoting 40S degradation, as reported previously ^5,11,12^. We note that the frequency of uridylation is likely underestimated: because RIOK3-bound 18S rRNAs are decay intermediates that undergo progressive degradation ^11^, they generate multiple sequencing reads per rRNA molecule, whereas rRNAs in the input fraction are largely intact and thus contribute fewer reads per molecule.

Given that RIOK3 kinase function is critical for eS26 release (Figures 1B and S1G), we next asked whether it is required for uridylation by comparing uridylated reads distributions between RIOK3 WT and ΔKD immunoprecipitates. Consistent with our *in vitro* biochemical characterization (Figures 1B and 1C), RIOK3 ΔKD-bound 18S rRNA exhibited markedly reduced uridylation levels (Figures S2D and 2E). To enable robust comparison across conditions, we calculated cumulative distributions of untemplated oligo-U lengths, a metric that is less sensitive to differences in sequencing depth. Again, we observed a pronounced reduction in oligo-U tails in the RIOK3 ΔKD samples (Figure 2F).

Since nonfunctional 18S rRNA decay undergoes decay via the GCN2-RNF10-RIOK3 axis ^5^, we next purified decoding-incompetent 40S subunits and sequenced the associated 18S rRNA decay intermediates to test whether uridylated intermediates are present. Motif analysis of untemplated reads from these decoding-incompetent 18S molecules revealed similar oligo-U tracts at the 3’end of decay intermediates (Figure S2E), with higher levels of uridylation compared to 18S:WT counterpart (Figure 2G, black and red lines). Importantly, the abundance of uridylated 18S:A1824C rRNA was markedly decreased in GCN2 KO, RNF10 KO, and RIOK3 KO cells (Figure 2G), consistent with the role of this cascade in 18S NRD ^5^. Together, these findings indicate that RIOK3 promotes uridylation of 18S rRNA during 40S turnover.

### TUTases mediate uridylation of 18S rRNA during decay

Given the oligo-U signature found in RIOK3-bound and nonfunctional 18S rRNA decay intermediates (Figures 2B and S2E) and prior studies implicating terminal uridylyltransferases in the turnover of structured noncoding RNAs ^13,19,24,31^, we next tested their roles in uridylating 18S rRNA. We first confirmed that the TUT4 KO, TUT7 KO, and TUT4/7 double-knockout (DKO) did not affect uS3 ubiquitination upon HTN treatment (Figure S2F). We then sequenced RIOK3-bound 18S rRNA decay intermediates from TUT7 and TUT4/7 DKO cells. Analysis of oligo-U distributions of RIOK3-bound 18S rRNA decay intermediates in TUT7 KO cells showed a moderate reduction in uridylation (Figure 2H), consistent with its partial redundancy in uridylating miRNAs ^23,32^. Notably, TUT4/7 DKO resulted in a more pronounced decrease in uridylation levels, particularly among reads containing more than five uridines (Figure 2H). Despite the overall reduction in uridylation, motif analysis of untemplated reads from TUT4/7 DKO cells still revealed enrichment of oligo-U tracts within 18S rRNA decay intermediates (Figure S2G). Consistent with this notion, decoding-incompetent 18S rRNA was not stabilized in TUT4/7 DKO compared to parental cells (Figures S2H and S2I). Together, these results support a model in which TUT4 and TUT7 are the primary enzymes driving uridylation, while additional factors may contribute to residual levels of uridylation, which appears sufficient to promote 18S rRNA decay. Notably, this conclusion is consistent with the partially redundant, but critical, role of the TUTases in uridylating miRNAs^23,32^.

### DIS3L2 degrades uridylated 18S rRNA during 40S subunit turnover

Oligo-uridylation frequently serves as a signal for RNA degradation by the 3ʹ-5ʹ exonuclease DIS3L2^19^, which has been shown to target a diverse range of uridylated RNA species, including tRNAs, pre- and mature miRNAs, and 5.8S rRNA biogenesis intermediates ^13,23,24,33–35^. To determine whether DIS3L2 participates in degrading uridylated 18S rRNA, we first assessed the stability of the decoding-incompetent 18S:A1824C rRNA in DIS3L2 KO cells. To this end, 18S:A1824C rRNA was constitutively expressed in DIS3L2 KO cells and its abundance was monitored over a 24-hour time course following inhibition of RNA polymerase I transcription. In parental cells, the mutant 18S rRNA was rapidly degraded, exhibiting a half-life of approximately 4.1 ± 1.7 hours, whereas 18S:WT rRNA remained stable for more than 24 hours (Figure 3A) as previously reported ^5,36^. Strikingly, loss of DIS3L2 markedly stabilized the mutant 18S rRNA, extending its half-life beyond 24 hours. Consistent with this finding, steady-state levels of the mutant 18S rRNA were increased in DIS3L2 KO cells. This stabilization phenotype was rescued by over-expression of DIS3L2 WT but not a catalytic mutant (D391N) (Figures 3B-3C).

**Figure 3.**
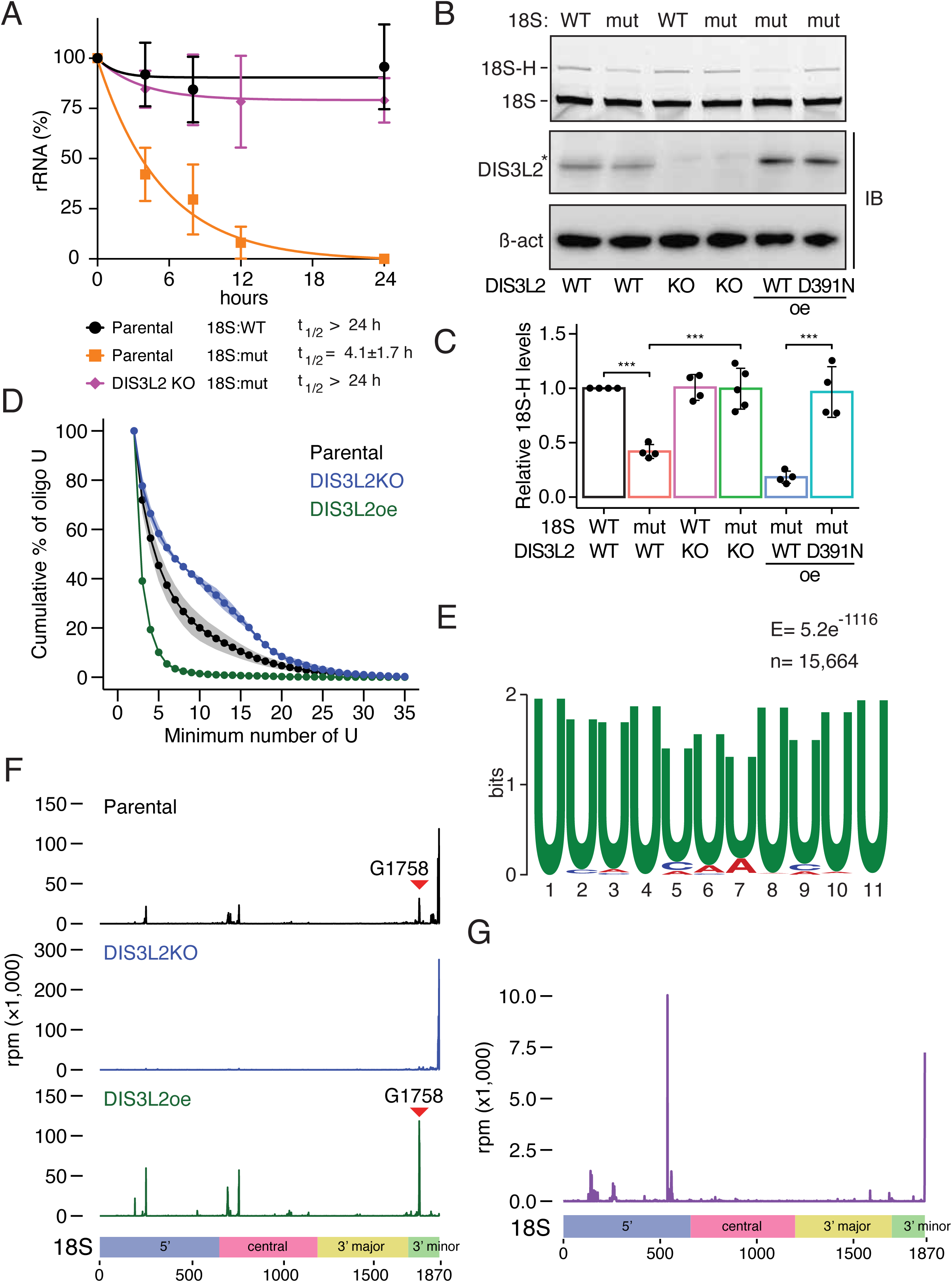
DIS3L2 degrades uridylated 18S rRNA. (A) Half-life measurements of 18S:A1824C (mut) in parental (orange) and DIS3L2 KO (purple) backgrounds compared to 18S:WT in parental cells (black). Half-lives are denoted with ± SD. (B) Primer extension analysis of 18S NRD reporters in HEK293T parental or DIS3L2 KO cells complemented with WT or catalytically inactive DIS3L2 (D391N). mut: 18S:A1824C. Bottom: immunoblots for DIS3L2. Asterisk denotes a non-specific band. (C) Quantification of the primer extension assays shown in (B) (n≥ 4). Student’s t-test is indicated by asterisks. *: p < 0.05, **: p < 0.01, and ***: p < 0.001 (D) Cumulative percentage of uridylated reads across the 18S rRNA from RIOK3-bound 18S rRNA isolated from HTN-treated HEK293T parental (black), DIS3L2 KO (blue), and DIS3L2 overexpression (oe) cells (green), (n≥ 2) (E) MEME results from untemplated nucleotides at 3’ ends of 18S decay intermediates isolated from DIS3L2 KO cells. E represents the E-value significance; n denotes the number of contributing sites out of 22,631 input sequences. (F) Representative gene models of 18S rRNA illustrating the distribution of 3ʹ ends of RIOK3-bound 18S rRNA isolated from HEK293T parental (black), DIS3L2 KO (blue), and DIS3L2 oe cells (green), (n≥ 2). Red arrowheads indicate enriched truncation in DIS3L2 oe cells in the 3’ minor domain. (G) Representative gene model of 18S rRNA illustrating the 3ʹ ends of DIS3L2 D391N-crosslinked reads. Enrichment of uridylated reads across multiple domains of the 18S rRNA indicates that DIS3L2 engages not only 3’ ends but also internal decay fragments. Data were obtained from GEO (accession number: GSE81537) ^13^.

We next examined uridylation levels of RIOK3-bound 18S rRNA isolated from DIS3L2 KO cells treated with HTN (Figure S3A) by Nanopore direct RNA sequencing. Loss of DIS3L2 led to a marked accumulation of uridylated reads compared to parental cells (Figure 3D). Motif analysis of untemplated reads from DIS3L2 KO cells revealed an even more striking enrichment of longer oligo-uridine tracts (Figure 3E) relative to those found in parental cells (Figure 2B). Conversely, overexpression of DIS3L2 largely diminished the accumulation of these uridylated species (Figure 3D).

Given the 3’ to 5’ end directionality of Nanopore direct RNA sequencing, the 3’ ends of the reads correspond to rRNAs truncated from their 3’ termini. Mapping the 3’ ends of all aligned reads, we observed distinct peaks across the 5’, central, and 3’ minor domains of 18S rRNA in parental cells (Figure 3F, black trace), consistent with progressive endo- and exoribonucleolytic cleavages during decay ^11^. In contrast, DIS3L2 KO samples displayed a dominant peak at the 3’ end, accompanied by a marked loss of internal cleavage signatures (Figure 3F, blue trace), indicative of full-length 18S rRNA. Importantly, although uridylation levels were elevated in DIS3L2 KO cells (Figure 3D), these uridylated reads were predominantly positioned at the 3’ termini of intact 18S rRNA (Figure S3B, blue trace), consistent with stabilization and accumulation of terminally uridylated 18S rRNA rather than increased degradation. Intriguingly, the loss of internal endoribonucleolytic cleavages in DIS3L2 KO cells suggests that DIS3L2-mediated exoribonucleolytic decay primes subsequent endoribonucleolytic decay processes. In contrast, overexpression of DIS3L2 (DIS3L2oe) enhanced decay signatures, characterized by increased peak intensity across multiple domains (Figure 3F, green trace). Strikingly, the peak at the very 3’ end of 18S rRNA was completely diminished in DIS3L2oe cells, with the appearance of a new peak at G1758 (Figure 3F, green trace, red arrowhead), indicating enhanced degradation of 18S decay intermediates.

These observations were further supported by analysis of read coverage profiles, which report sequencing depth across 18S rRNA. RIOK3-bound 18S rRNA in parental cells showed significant reductions in coverage corresponding to regions that undergo degradation (Figure S3C, black). DIS3L2 KO samples exhibited a gradual decline in read coverage toward the 5’ end of 18S rRNA (Figure S3C, blue trace), lacking sharp drop-offs in coverage seen in parental cells. Conversely, DIS3L2 overexpression resulted in a pronounced loss of coverage in the central and 3’ minor domains (Figure S3C, green), indicative of removal approximately half of the 3’ minor domain upon DIS3L2 overexpression (Figure S3D). Together, these findings establish DIS3L2 as the key exoribonuclease that triggers degradation downstream of 18S rRNA uridylation. Overall, these data reveal critical steps in ribosome quality surveillance that couple 18S rRNA uridylation to 40S subunit turnover.

### DIS3L2 acts at multiple steps of 18S rRNA degradation

To dissect whether DIS3L2 acts at other steps in 18S rRNA decay, we re-analyzed DIS3L2 enhanced UV cross-linking and immunoprecipitation (eCLIP) data generated with the catalytically inactive mutant D391N, which allows stabilization and capture of otherwise transient decay substrates ^13^. Analysis of DIS3L2 D391N-crosslinked 18S rRNA fragments revealed uridylated reads at the 3’ end and in the 5’ domain of 18S rRNA (Figure 3G). Notably, the enrichment of uridylated species just upstream of endoribonucleolytic cleavage sites in the 5’ domain suggests that TUTases act on decay intermediates following endoribonucleolytic cleavage, priming these fragments for downstream degradation. Further supporting the role of DIS3L2 in degrading 18S rRNA, we found it was present in the free RNP and 40S fractions in unstressed cells. Upon HTN treatment, DIS3L2 was further enriched in the 40S fraction (Figure S3E). Given that RIOK3 binds and remodels ubiquitinated 40S subunits to license decay, these findings support a model in which further uridylation occurs possibly after RIOK3 dissociation, thereby ensuring efficient clearance of 18S rRNA by DIS3L2. Collectively, this observation suggests that DIS3L2 acts on both the 3’ end of 18S rRNA and uridylated decay intermediates generated by endoribonucleolytic cleavage to complete 40S subunit turnover.

### *In vitro* reconstitution of uridylation-dependent 18S rRNA decay

To define the biochemical basis of uridylation-dependent 18S rRNA decay, we reconstituted the reaction *in vitro* using purified recombinant TUT7 catalytic domain (CD) and DIS3L2 (Figure S4A). We then utilized the amplicon-based sequencing approach described above (Figure S2A) to quantify uridylation levels at the 3’ end of 18S rRNA. Incubation of ubiquitinated 40S ribosomal subunits with TUT7CD yielded short uridine extensions relative to ubiquitinated 40S subunits alone (Figure 4A, pink trace). In contrast, addition of RIOK3 dramatically increased the length and abundance of uridylated reads (Figure 4A, blue trace). Importantly, this stimulation required RIOK3 kinase function, as RIOK3 ΔKD failed to promote robust uridylation (Figure 4B).

**Figure 4.**
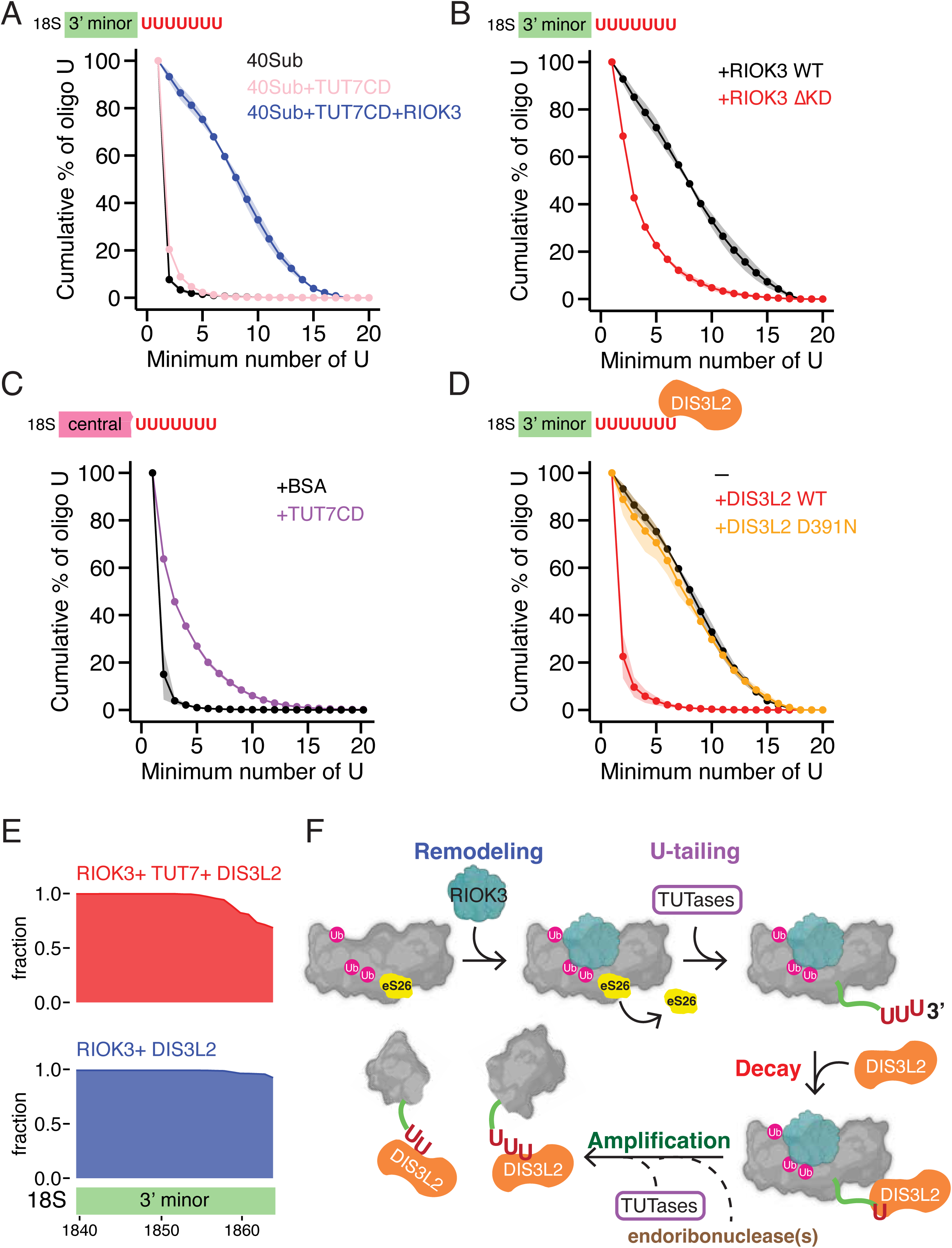
*In vitro* reconstitution of 18S rRNA decay. (A) Cumulative percentage of uridylated reads at the 3’ end of 18S rRNA in reactions containing ubiquitinated 40S (40Sub, black), 40Sub +TUT7CD (pink), or 40Sub + TUT7CD + RIOK3 (blue), (n≥ 2). Schematic depicts uridylation at the 3’ minor domain (top). (B) Similar to (A), cumulative percentage of uridylated reads at the 3’ end of 18S rRNA in reactions containing 40Sub, TUT7CD, and RIOK3 WT (black) or RIOK3 ΔKD (red), (n≥ 2). (C) Cumulative percentage of uridylated reads derived from the central domain of 18S decay intermediates in reactions containing BSA (black) or TUT7CD (purple), (n= 2). RIOK3-bound 18S decay intermediates isolated from HTN-treated TUT4/7 DKO cells were used as substrates for *in vitro* uridylation assays. See Figure S4D for corresponding uridylation levels in the 3’ minor domain. (D) Cumulative percentage of uridylated reads at the 3’ end of 18S rRNA in reactions containing 40Sub, TUT7CD, and RIOK3 with DIS3L2 WT (red) or the catalytic mutant DIS3L2-D391N (orange) or without DIS3L2 (black), n≥ 2. See also Figure S4E for degradation extending upstream within the 3’ minor domain. (E) Representative read coverage of 18S 3’ minor domain in reactions containing 40Sub, RIOK3, and DIS3L2 WT with (red) or without TUT7CD-mediated uridylation (blue), (n= 2). Decreased density at the 3’ end indicates exoribonucleolytic degradation extending upstream from the uridylated termini. (F) Working model for uridylation- and DIS3L2-mediated 40S turnover. RIOK3 recognizes ubiquitinated 40S ribosomes and remodels the subunit by displacing ribosomal protein eS26, thereby exposing the 3ʹ end of 18S rRNA. This remodeling step licenses terminal uridylation by TUTases, which add oligo-uridine tails to the 3ʹ end of 18S rRNA. DIS3L2 engages uridylated 18S rRNA to initiate 3ʹ-5ʹ exoribonucleolytic decay. 3’-end degradation promotes endoribonucleolytic cleavage, generating intermediates for further uridylation and DIS3L2 engagement.

Given that RIOK3 releases eS26 from ubiquitinated 40S subunits (Figure 1), we reasoned that eviction of eS26 exposes the 3’ ends of 18S rRNA, thereby allowing TUTases to catalyze uridylation. To test this idea further, we added recombinant PNO1, which covers the 3’ end of 18S rRNA in a manner similar to eS26 ^11,37,38^ (Figure S4B), and assessed TUT7CD-mediated uridylation. Indeed, adding PNO1 reduced uridylation levels (Figure S4C). Collectively, our results indicate that RIOK3-mediated release of eS26 renders the 3’ termini of 18S rRNA accessible to TUTases for uridylation.

Our Nanopore RNA sequencing results indicate that RIOK3-bound 18S rRNA is endoribonucleolytically cleaved within the 5’ and central domains in cells (Figure 2D). To corroborate this observation, we isolated RIOK3-bound 18S rRNA decay intermediates from HTN-treated TUT4/7 DKO cells, in which uridylation is reduced (Figure 2H) and the 18S rRNA is truncated within the 5’, central, and 3’ minor domains, and used these intermediates as substrates for TUT7CD. *In vitro* uridylation assays showed that TUT7CD efficiently uridylated the 3’ ends of these decay intermediates derived from both the central domain (Figure 4C) as well as 3’ minor domain (Figure S4D), supporting the notion that TUTases act on 18S decay intermediates after endoribonucleolytic cleavages.

Next, we tested whether uridylated 18S rRNA could be further degraded by DIS3L2 *in vitro*. Recombinant DIS3L2 was added to uridylation reactions catalyzed by recombinant RIOK3 and TUT7CD. Addition of DIS3L2 WT efficiently shortened uridine tails in the 3’ end of 18S rRNA, whereas the catalytic mutant D391N failed to do so (Figure 4D). In addition, we observed that DIS3L2 WT promotes 18S rRNA degradation extending upstream into the 3’ minor domain (Figure S4E). Importantly, DIS3L2 failed to degrade 18S rRNA in the absence of TUT7CD (Figure 4E), indicating uridylation in the 3’ end of 18S rRNA is required for DIS3L2-depedent decay. Together, these findings define a pathway for mammalian 18S rRNA quality control in which RIOK3 drives remodeling of ubiquitinated 40S subunits to enable TUTase-mediated uridylation of 18S rRNA. The resulting U-tails serve as a degradation signal for the 3ʹ-5ʹ exoribonuclease DIS3L2. This stepwise mechanism couples ribosome remodeling with rRNA decay, establishing uridylation-dependent DIS3L2 activity as the terminal execution step of 40S subunit turnover.

## DISCUSSION

Selective turnover of 40S ribosomal subunits is triggered by diverse stresses that cause impaired translation. Although the upstream triggers differ— including decoding-incompetent 18S rRNA, nutrient stress, disrupted biogenesis of the 60S ribosomal subunits, and ribosome-targeting drugs— the downstream players have very recently been shown to be strikingly similar. Still, the turnover mechanism that clears 40S subunits previously remained unclear.

Here, we define the mechanism by which these ubiquitinated 40S subunits are committed to decay. Our data support a pathway in which ubiquitinated 40S subunits are remodeled to expose the 3ʹ terminus of 18S rRNA, which is then uridylated by TUTases and degraded by DIS3L2 (Figure 4F). Importantly, our work identifies the remodeling step that converts ubiquitinated 40S ribosomal subunits, which are still intact ribonucleoprotein particles, into degradable RNA substrates. Hence, this study provides a direct mechanistic link between early translation perturbation and selective ribosomal subunit turnover.

Loss of DIS3L2 stabilizes the decoding-incompetent 18S rRNA and leads to accumulation of uridylated 18S rRNA species, whereas DIS3L2 overexpression exhibits the opposite effect (Figures 3A-3E). DIS3L2 degrades uridylated 18S rRNA in a 3’-5’ manner, generating a discrete decay intermediate with a defined 3’ terminus at approximately G1758, particularly in DIS3L2oe cells (Figure 3F, red arrowhead). This truncation removes two of the three critical monitoring bases ^39–42^ (A1824 and A1825) in the decoding center (Figure S3D), effectively rendering the 40S subunit translationally inactive. Together, these findings identify DIS3L2 as the 3’-5’ exoribonuclease that executes 18S rRNA degradation during 40S subunit turnover. Bacterial 16S rRNA quality control has been linked to RNase R-dependent pathways ^43,44^, and DIS3L2 belongs to the same RNase II/RNB exoribonuclease family ^34,45,46^. This commonality implies that a related decay pathway targets small ribosomal subunits even though distinct upstream signals commit ribosomal RNAs to degradation.

Moreover, our Nanopore sequencing and DIS3L2 eCLIP data ^13^ further indicate that DIS3L2 functions at multiple stages of the 18S rRNA decay process, acting both early in initiating degradation and later in clearing terminal fragments. First, DIS3L2 KO leads to the accumulation of uridylated reads at the 3ʹ termini of 18S rRNA together with a loss of internal decay signatures (Figure 3F). This pattern argues that DIS3L2-mediated 3ʹ-5ʹ exoribonucleolytic decay promotes the broader degradation program, potentially priming substrates for subsequent endoribonucleolytic events. While RNase R rearranges bacterial 30S subunits during 16S rRNA degradation ^44^, it is plausible that DIS3L2 similarly remodels human 40S subunits to induce endoribonucleolytic cleavage. Second, crosslinking analysis reveals that the catalytic mutant DIS3L2-D391N binds uridylated 18S rRNA decay intermediates derived from the 5’ domain (Figure 3G), indicating that DIS3L2 also engages internal decay fragments once they become uridylated (Figure 4F). In this model, initial DIS3L2-mediated trimming from the 3’ termini may promote endoribonucleolytic cleavages, after which the resulting fragments are further uridylated by TUTases and re-engaged by DIS3L2 for progressive degradation. Future structural and biophysical studies of DIS3L2-mediated 40S subunit decay will provide spatially and temporally resolved insights into how uridylated 18S rRNA is recognized and degraded within remodeled 40S subunits.

Together, our findings define the downstream mechanism by which mammalian 40S ribosomal subunits are committed to destruction. 40S ribosomal particles targeted for turnover are remodeled to expose the 3ʹ end of 18S rRNA, marked by terminal uridylation, and then eliminated by DIS3L2. This pathway reveals how ribosome remodeling, terminal RNA modification, and exonucleolytic decay are coupled during mammalian ribosome quality control.

## ACKNOWLEDGMENTS

We would like to thank Jeff Carrell and Megan Karwan at the NCI-Frederick Flow Cytometry Core Facility for assistance in cell sorting; Ronald Holewinski and Thorkell Andresson at the NCI-Frederick Protein Characterization Laboratory for mass spectrometric analysis. This work was supported by the Intramural Research Program of the National Institutes of Health, National Cancer Institute, Center for Cancer Research (1ZIABC012037 to C.W. and 1ZIABC011566 to S.G.).

## AUTHOR CONTRIBUTIONS

Conceptualization, A.S. and C.C.C.W.; Methodology, A.S. and A.R.C.; Investigation, A.S., A.R.C., A.S., J.T.M., and C.C.C.W.; Resources, A.R.C., E.O., A. Y., and J.T.M.; Visualization, A.S., C.C.C.W., and W.G.; Data Curation, W.G.; Writing, A.S. and C.C.C.W.; Supervision, S. G. and C.C.C.W.

## DATA AVAILABILITY

Sequencing data generated in this study have been deposited in the SRA database under project number: PRJNA1451181.

## EXPERIMENTAL PROCEDURES

### Cell lines and culture conditions

HEK293T and Flp-In T-REx HeLa cells were grown in Dulbecco’s modified medium (DMEM) supplemented with 10% fetal bovine serum, 2 mM *L*-glutamine. HEK293T parental, TUT7 KO, TUT4/7 KO, and DIS3L2 KO cells were generated previously^22,31^. Cells were maintained in a 5% CO2 humidified incubator and passaged every 2-3 days.

### Generation of knockout cell lines

sgRNAs targeting human 3’-5’ exoribonuclease DIS3L2 (5’-CACCGTGGATAAAGTGGCTGCCGAG-3’ and 5’-AAACCTCGGCAGCCACTTTATCCAC-3’) were annealed and cloned into lentiCRISPRv2-puro vector (Addgene, #98290) and lentiviral particles were generated as previously described ^4^. 1 mL of lentiCRISPRv2 viral particles were incubated with 2 x 10^5^ Flp-In T-REx HeLa cells and polybrene transduction reagent (10 μg/mL) (Sigma, TR-1003) at 37°C for 30 minutes. This mixture was then plated onto a 6-well plate. 24 hours after transduction, media was changed to DMEM supplemented with 10% FBS and 2 mM *L*-glutamine. 48 hours post transduction, cells were selected with puromycin (1 μg/mL) (P8833, Sigma). Monoclonal DIS3L2 KO cell lines were generated using a limiting dilution method. Loss of DIS3L2 was validated by immunoblotting.

### Generation of 18S-H expressing cell lines

2 x 10^5^ DIS3L2 KO HeLa cells were seeded in a 6-well plate a day prior to transfection with 1 μg of either 18S-H:WT or 18S-H:A1824C plasmid and 1 μg of PiggyBac transposase (System Biosciences) as previously described ^4^. 4 days after transfection, cells were trypsinized for single cell sorting. tagBFP-positive cells were sorted into three 96-well plates. Expression of 18S-H rRNA was subsequently confirmed by primer extension analysis.

### *In vitro* ubiquitination assay

Ribosomes were isolated from whole-cell extracts by layering the extract onto a 35% sucrose cushion followed by centrifugation at 100,000 × g for 1 h. The ribosome pellet was resuspended in ribosome resuspension buffer (50 mM Tris-HCl [pH 8.0], 25 mM KCl, and 5 mM MgCl2). 20 U of ribosomes were ubiquitinated *in vitro* using a ubiquitination kit (Enzo Life Sciences, BML-UW9920) according to the manufacturer’s protocol. Briefly, the reaction contained 1× ubiquitination buffer, IPP (100 U/mL), 1 mM DTT, 5 mM Mg-ATP, 100 nM E1 enzyme, 500 nM E2 enzyme, 450 nM RNF10, 2.5 μM biotinylated ubiquitin, and 2 U SUPERase-In RNase inhibitor. Reactions were carried out at 37°C for 3 h.

### *In vitro* eS26 displacement assay

#### Preparation of 40S ribosomal subunits

Frozen cell pellets collected from four 15-cm dishes of Flp-In T-REx HeLa USP10 RIOK3 DKO uS3-HA cells were lysed in 1 mL lysis buffer per plate (30 mM HEPES [pH 7.5], 50 mM KCl, 10 mM MgCl2, 220 mM sucrose, 2 mM DTT, 0.5 mM EDTA, 0.5% Igepal, and 0.5 mg/mL heparin, 1× cOmplete Mini Protease Inhibitor Cocktail [Roche, 11836170001], 20 U SUPERaseIn). Lysates were incubated on ice for 10 min, centrifuged at 10,000 × g for 5 min at 4°C, and the supernatants were transferred to fresh tubes. The lysates were further clarified by centrifugation at 21,000 × g for 30 min at 4°C. PEG 20,000 was added to the clarified lysate to a final concentration of 1.35%, followed by incubation on ice for 15 min. After centrifugation at 20,000 × g for 12 min at 4°C, the supernatant was collected, adjusted to 130 mM KCl, and incubated for 5 min. PEG 20,000 was then increased to a final concentration of 4.5%, and samples were incubated on ice for an additional 10 min. Ribosomal particles were collected by centrifugation at 17,500 × g for 10 min at 4°C, and the resulting pellet was resuspended in buffer R (30 mM HEPES [pH 7.5], 125 mM KCl, 7.5 mM MgCl2, 2 mM DTT, and 0.5 mg/mL heparin, 1× cOmplete Mini Protease Inhibitor Cocktail). Resuspended ribosomes were loaded onto 10-30% sucrose gradients prepared in high-salt gradient buffer (25 mM HEPES [pH 7.5], 550 mM KCl, 5 mM MgCl2, and 2 mM DTT) and centrifuged at 39,000 rpm for 3 h at 4°C in an SW40 rotor (Beckman Coulter). Gradients were fractionated into 1 mL fractions, and fractions corresponding to the 40S peak were pooled. Pooled 40S fractions were then incubated overnight with washed anti-HA beads to capture uS3-HA-containing 40S ribosomes.

#### *In vitro* ubiquitination of immunoprecipitated uS3-HA-containing 40S ribosomes

Anti-HA beads bound to uS3-HA-containing 40S ribosomes were washed once with IP lysis buffer (50 mM HEPES [pH 7.5], 5 mM MgCl₂, 150 mM NaCl, 1% Triton X-100, 1× cOmplete Mini Protease Inhibitor Cocktail) and once with ribosome resuspension buffer (50 mM Tris-HCl [pH 8.0], 25 mM KCl, 5 mM MgCl₂). Bead-bound 40S ribosomes were then subjected to in vitro ubiquitination reaction as described previously. Following the ubiquitination reaction, beads were washed once with ribosome resuspension buffer and resuspended in the same buffer for subsequent reactions.

#### RIOK3-mediated eS26 displacement assay

Ubiquitinated uS3-HA-bound 40S ribosomes immobilized on anti-HA beads were incubated with 1.6 μM recombinant RIOK3 WT, RIOK3-ΔKD, or BSA control, in a reaction containing 20 mM Tris-HCl [pH 8.0], 150 mM KCl, 20 mM MgCl2, 1 mM ATP, and 20 U SuperRaseIn. Reactions were incubated for 1 h at 37°C. Following incubation, supernatants were collected and mixed with 1× SDS sample buffer. Beads were washed once with ribosome resuspension buffer, and bead-associated material was recovered in 1× SDS sample buffer.

### RIOK3 immunoprecipitation

For analysis of RIOK3-associated 18S rRNA, cells grown in a 15 cm dish were transfected with 20 μg of pcDNA-RIOK3-FLAG plasmid. 48 h post transfection, cells were treated with 3 μg/mL HTN for 30 mins and lysed in IP lysis buffer (50 mM HEPES [pH 7.5], 150 mM NaCl, 5 mM MgCl2, 0.5% Triton X-100, 1× cOmplete Mini Protease Inhibitor Cocktail [Roche, 11836170001], 200 U SUPERase-In RNase inhibitor, and 400 U murine RNase inhibitor [New England Biolabs, M0314]). Lysates were clarified and incubated with 50 μL anti-DYKDDDDK magnetic agarose beads (Thermo Fisher Scientific, A36797) with rotation at 4 °C for 16 h. Beads were subsequently washed three times with IP wash buffer (50 mM HEPES [pH 7.5], 150 mM NaCl, 5 mM MgCl2, 1% Triton X-100, 0.5 mM DTT). FLAG-tagged RIOK3 complexes were eluted using FLAG peptide (0.5 mg/mL in 100 μL PBS supplemented with 60 U SUPERase-In RNase inhibitor) for 30 min at 4 °C. RNA was then isolated from the eluate by phenol-chloroform extraction and resuspended in 10 μL RNase/DNase-free water.

### *In vitro* uridylation assay

40S ribosomal subunits were prepared as described above from Flp-In T-REx HeLa USP10/RIOK3 DKO uS3-HA cells treated with HTN (3 μg/mL) for 30 min before 40S isolation. 46 nM purified 40S subunits were incubated in reactions containing 20 mM Tris-HCl [pH 8], 150 mM KCl, 20 mM MgCl2, 0.5 mM DTT, 960 nM recombinant RIOK3 WT, 1 mM ATP, 0.25 mM UTP, 20 U SUPERaseIn, and 560 nM recombinant TUT7CD at 37°C for 2 h. For DIS3L2 *in vitro* assays, recombinant DIS3L2 WT or D391N was added to the above reactions at 205 nM and incubate at 37°C for 30 min. In experiments examining the effect of PNO1, 46 nM purified 40S subunits were incubated with 1.8 µM RIOK3 for 1 h, followed by addition of 1.7 µM PNO1 or BSA and incubation for 30 min; 800 nM TUT7CD and 0.25 mM UTP were then added, and reactions were incubated for an additional 1 h. For in vitro uridylation of 18S rRNA in the central domain, RIOK3-bound 18S decay intermediates were isolated from ∼20 million HTN-treated TUT4/7DKO cells as described above. RIOK3-IP eluates were incubated in reactions containing 20 mM Tris-HCl [pH 8], 150 mM KCl, 20 mM MgCl2, 0.5 mM DTT, 0.25 mM UTP, 20 U SUPERaseIn, and 560 nM BSA or recombinant TUT7CD at 37°C for 2 h.

RNA was isolated by phenol-chloroform extraction followed by sodium acetate/isopropanol precipitation. Recovered RNA was resuspended in 10 mM Tris-HCl [pH 8.0], heated to 80°C for 2 min, chilled on ice, and subjected to 3ʹ dephosphorylation using T4 polynucleotide kinase for 1 h at 37°C. A pre-adenylated 3ʹ linker (5Phos/App-NNNNNNCACTCGGGCACCAAGGA/3ddC) was then ligated using T4 RNA ligase 2 truncated and incubating for 3 h at 37°C, followed by overnight sodium acetate precipitation at -80°C. The ligated RNA was resuspended in water and reverse transcribed using SuperScript III at 55°C for 30 min. RNA templates were hydrolyzed with NaOH, and cDNA was recovered by sodium acetate/isopropanol precipitation. cDNA was amplified by an initial PCR using NEBNext Ultra II Q5 Master Mix (New England Biolabs, M0544X), reverse primer (5’-TCCTTGGTGCCCGAGTG-3’) and forward primer (5’-CCCTACACGACGCTCTTCCGATCTNNNNNTTTCCGTAGGTGAACC-3’) for the 3’ minor domain or (5’-CCCTACACGACGCTCTTCCGATCTNNNNNCTCCAATAGCGTATATTAAAG-3’) for the central domain, purified with AMPure XP beads, and subjected to a second indexing PCR with barcoded primers. Barcoded libraries were resolved on 8% native PAGE, extracted using sodium acetate/isopropanol precipitation, and on an Illumina NextSeq 1000/2000.

3’ adapter (NNNNNNCACTCGGGCACCAAGGA) was trimmed off using skewer ^46^. Trimmed reads were aligned to human 18S rRNA using STAR ^47^. For counting untemplated U reads, only sequences beginning with “TT” were considered. Thymidines were counted, while non-T nucleotides (A, C, G) were treated as mismatches and tolerated up to 3 mismatches. Mismatches immediately followed by a thymidine were permitted, and scanning was terminated when the mismatch threshold was exceeded. Same criteria were applied to reanalysis of DIS3L2 D391N-corsslinked reads (Figure 3G) and data were obtained from GEO (accession number: GSE81537)^13^. Cumulative percentage of uridylation was calculated as fraction of reads containing at least the indicated number of untemplated uridines and plotted in Figures 4 and S4.

### Library preparation for Nanopore sequencing

For nanopore sequencing library preparation, purified RIOK3-associated RNA was polyadenylated using *E. coli* poly(A) polymerase (New England Biolabs, M0276S) according to Oxford Nanopore protocols. The poly(A)-tailed RNA was then used to generate sequencing libraries using a Direct RNA Sequencing Kit (Oxford Nanopore, SQK-RNA004) following the manufacturer’s recommendations. Libraries were loaded onto SpotON flow cells and sequenced on a GridION instrument.

Basecalling was performed by the MinKNOW software. Reads were mapped to 18S rRNA reference sequence using minimap2 ^48^ with the following parameters: ‘‘–secondary=no’’. Aligned reads were analyzed using custom scripts written in Python 3 and R ^49^. Oligo-U counting is as described above.

### Library preparation for Illumina sequencing

Libraries were prepared as described above in ***In vitro* uridylation assay.** For sequence motif analyses, untemplated reads starting with ‘ACG’ were removed to exclude biogenesis intermediates ^10,50^, and the resulting fasta files were analyzed by MEME ^51^.

### Co-immunoprecipitation analysis

Cells were transfected with pcDNA-RIOK3-FLAG plasmid for 48 h. At 48 h post-transfection, cells were treated with 3 μg/mL HTN for 3 h and lysed in IP lysis buffer (50 mM HEPES [pH 7.5], 5 mM MgCl₂, 150 mM NaCl, 1% Triton X-100, 1× cOmplete Mini Protease Inhibitor Cocktail). For crosslinking, lysates were treated with 1 mM dithiobis (succinimidyl propionate) for 10 min at room temperature. The reaction was quenched by addition of 200 mM Tris-HCl [pH 7.5]. Lysates were then clarified by centrifugation at 14,000 × g for 10 min. Clarified extracts were incubated with 30 μL of either anti-FLAG beads or IgG control beads for 3 h, followed by five washes with IP wash buffer (50 mM HEPES [pH 7.5], 5 mM MgCl₂, 300 mM NaCl, and 1.5% Triton X-100). Bound proteins were eluted in 30 μL of 2× SDS sample buffer and immunoblotted.

### Polysome profiling

Cells were treated with HTN (3 μg/mL) for 30 min prior to lysis in footprint lysis buffer (20 mM Tris-HCl [pH 8.0], 150 mM KCl, 15 mM MgCl2, 1% Triton X-100, 1 mM DTT, and 0.1 mg/mL cycloheximide). Lysates were clarified by centrifugation at 21,000 × g for 10 min and clarified lysate containing 500 μg of RNA was loaded onto a 10-30% sucrose gradient. Gradients were centrifuged in an SW40Ti rotor for 3 h at 4°C and fractionated using a Biocomp fractionator. Proteins in each fraction were precipitated with 8% trichloroacetic acid (TCA) and analyzed by immunoblotting.

### Purification of recombinant DIS3L2 proteins

6His-SUMO-DIS3L2 in pET32a was a gift from Sandra Wolin. Mutant D391N was created by Gibson assembly and verified by whole plasmid sequencing. The proteins were expressed in Rosetta (DE3) cells (Novagen) induced with 1 mM IPTG and grown overnight at 16 °C. Cells were lysed by sonication in 50 mM HEPES [pH 7.5], 150 mM NaCl and the clarified supernatant was incubated with NiNTA-agarose resin (Qiagen) equilibrated in 50 mM HEPES [pH 7.5], 150 mM NaCl at 4 °C with gentle rotation for 1 h. Resin was first washed with 10 column volumes 50 mM HEPES [pH 7.5], 150 mM NaCl, 30 mM Imidazole and then 10 column volumes 50 mM HEPES [pH 7.5], 500 mM NaCl, 30 mM Imidazole. Protein was eluted with 3 volumes of 50 mM HEPES [pH 7.5], 150 mM NaCl and 500 mM Imidazole. Fractions containing the protein were pooled and concentrated using an Amicon Ultra-15 10KDa MWCO (Sigma-Aldrich, UFC901008D). The SUMO tag was cleaved by adding 1/10 volume of 10X SUMO salt free buffer (500mM Tris-HCl [pH 8], 2% Igepal, 1.5M NaCl, 10mM DTT) and 6 units of SUMO protease (UlpI, ScienCell, MB9018) per microgram of full length tagged 6HSUMODIS3L2 for 16 h at 4°C. The reaction was diluted to 30mM imidazole (1:17) with 50mM HEPES [pH 7.5] and re-applied to NiNTA-agarose in 50 mM HEPES [pH 7.5], 150 mM NaCl. The flow through containing the cleaved DIS3L2 species was applied to a 1mL Hitrap heparin HP column (Cytiva) equilibrated with 10CV 25 mM HEPES [pH 7.5], 50 mM NaCl, 5 mM DTT, followed by a 10CV wash in 25 mM HEPES [pH 7.5], 50 mM NaCl, 5 mM DTT, and a 20 CV linear gradient from 50mM to 1M NaCl. Appropriate fractions were pooled and concentrated to 5 mL and loaded onto a size exclusion column (16/600 Superdex200pg, Cytiva) equilibrated with 50 mM HEPES [pH 7.5], 150 mM NaCl, 5 mM DTT. Fractions containing pure protein were pooled, concentrated, dialyzed overnight at 4°C against 20mM Tris-HCl [pH 8.0], 150 mM NaCl, 5 mM DTT and 20% glycerol, and stored at -80°C. The identities of the recombinant proteins were confirmed by mass spectrometry.

### Purification of recombinant TUT7CD proteins

The plasmid p6His-SUMO-TUT7-HA was created by Gibson assembly between the backbone of pSumo-His6-SUMO-AtTPR1(1-209) (Addgene #177858) and a truncated human TUT7 gene encoding amino acids 951 to 1495 with a C-terminal HA tag. The protein was expressed in Rosetta2 (DE3) cells with 1 mM IPTG, cells were lysed and purified by IMAC as above for DIS3L2. Concentrated 6HSUMOTUT7HA fractions were applied to a Superdex 200 size exclusion column equilibrated with 50 mM HEPES [pH 7.5], 150 mM NaCl, 1 mM DTT. Fractions containing pure protein were pooled, concentrated, dialyzed overnight at 4 °C against 20 mM Tris-HCl [pH 8.0], 150 mM NaCl, 1 mM DTT and 20% glycerol, and stored at -80°C.

### Purification of recombinant RIOK3 proteins

The plasmid p6His-SUMO-RIOK3 was created by Gibson assembly between the backbone of pSumo-His6-SUMO-AtTPR1(1-209) and either the full length human RIOK3 gene or a deletion of the kinase domain (residues 350 to 618). The proteins were expressed in Rosetta2 (DE3) cells with 1 mM IPTG, cells were lysed and purified by IMAC as above for DIS3L2. Concentrated fractions were applied to a Superdex 200 size exclusion column equilibrated with 50 mM HEPES [pH 7.5], 150 mM NaCl, 1 mM DTT. Fractions containing pure protein were pooled, concentrated, dialyzed overnight at 4°C against 20 mM Tris-HCl [pH 8.0], 150 mM NaCl, 1 mM DTT and 20% glycerol, and stored at -80°C.

The plasmid pMAL-c2G-RIOK3-MBP-6His was created by Gibson assembly between the backbone of pMAL-c2G (Addgene #75290) and either the full length human RIOK3 gene or a deletion of the kinase domain (residues 350 to 618) with a C-terminal 6X His tag. The proteins were expressed in Rosetta2 (DE3) cells with 1mM IPTG, cells were lysed, purified by IMAC followed by Superdex 200 size exclusion, concentrated and dialyzed as described above for p6His-SUMO-RIOK3.

### Purification of recombinant RNF10 protein

The plasmid pMAL-c2G-RNF10-6His was created by Gibson assembly between the backbone of pMAL-c2G and the human RNF10 gene with a C-terminal 6X His tag, replacing the MBP fusion protein. The protein was expressed in Rosetta2 (DE3) cells overnight at 18°C after induction with 1mM IPTG. Cells were lysed and the clarified supernatant was purified by IMAC as above for DIS3L2.

### Primer extension assay

Primer extension analysis was performed essentially as previous described ^4^. Total RNA was isolated from indicated cells using a Quick-RNA Miniprep Kit (Zymo, R1055) and quantified by NanoDrop 2000. Purified 1 µg RNA was incubated with 0.67 mM dNTPs and 0.67 mM Cy5-labeled oligonucleotide at 65 °C for 5 min and then placed on ice. Reverse transcription was performed with 5× First Strand Buffer and 100 U Maxima RT (Thermo Fisher Scientific, EP4701) at 53 °C for 30 min, followed by RNA hydrolysis with 0.1 N NaOH at 95 °C for 15 min. cDNA was precipitated, resolved on a 10% Criterion TBE-Urea polyacrylamide gel (Bio-Rad, 3450088), and imaged using an Amersham Typhoon scanner. Band intensities for endogenous 18S and 18S-H rRNA were quantified in ImageJ. Relative 18S-H levels were calculated as the ratio of 18S-H to endogenous 18S signal, and values from at least three biological replicates were used to determine the mean ± SD.

### Pulse-chase analysis

Pulse-chase analysis of 18S rRNA was performed essentially as described previously ^4^. Cells expressing 18S:WT or 18S:A1824C rRNA in parental or DIS3L2 KO backgrounds were seeded in 6-well plates 1 day before treatment. CX-5461 (Sigma, 5092650001) was added at 10 µM to inhibit Pol I transcription, and cells were harvested at the indicated time points for total RNA isolation using a Quick-RNA Miniprep Kit (Zymo, R1055). Samples were analyzed by primer extension, and relative tagged 18S rRNA levels were normalized to the 0 h time point, which was set to 100%.

### Statistical analysis

Significance was calculated using unpaired Student’s t-test (*: p < 0.05, **: p < 0.01, and ***: p < 0.001). Error bars denote the standard deviations.

